# Ultrasound-mediated drug diffusion, uptake, and cytotoxicity in glioblastoma 3D tumour sphere model

**DOI:** 10.1101/2023.10.26.564145

**Authors:** Janith Wanigasekara, Julie Rose Mae Mondala, Patrick J. Cullen, Brijesh Tiwari, James F. Curtin

## Abstract

A myriad of biological effects can be stimulated by the ultrasound (US) for the treatment of cancer. The objective of our research was to investigate the effect of ultrasound alone and in combination with the chemotherapeutic drugs such as doxorubicin (DOX) and temozolomide (TMZ) on human glioblastoma (GBM) and the human epidermoid carcinoma cancer 2D and 3D cell cultures. Results indicated that the US 96-probe device could induce tumour sphere cytotoxicity in a dosage- and time-dependent manner, with multiple treatments augmenting this cytotoxic effect. With enhanced cytotoxicity, US decreased tumour sphere growth metabolic activity, disrupted spheroid integrity, and heightened the occurrence of DNA double strand breaks, resulting in damage to tumour spheres and an inability to rebuild tumour spheres after multiple US treatments. The combination of US and TMZ / DOX enhanced the efficiency of treatment for GBM and epidermoid carcinoma by enhancing induced cytotoxicity in 3D tumour spheres compared to 2D monolayer cells, and also by increasing the incubation time, which is the most crucial way to differentiate between the effectiveness of drug treatment with and without US. In conclusion, our data demonstrate that US enhances drug diffusion, uptake, and cytotoxicity using 3D spheroid models when compared with 2D cultures. It also demonstrates the significance of 3D cell culture models in drug delivery and discovery research.

**Graphical Abstract:** 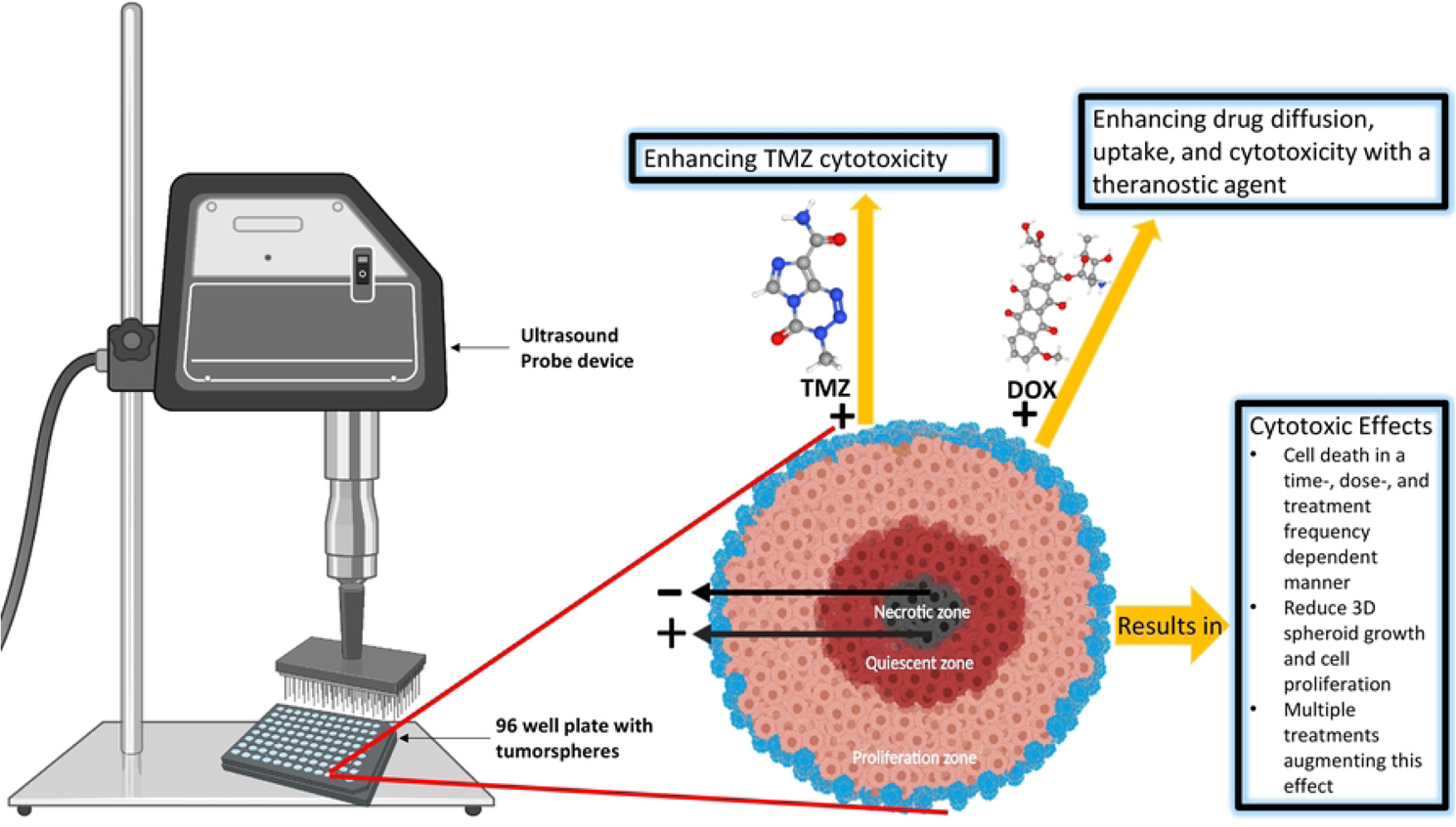

## 1. Introduction

Glioblastoma (GBM) is a grade IV glioma in adults, characterized as IDH wild type, according to the fifth edition of the WHO classification and defined from a histology and molecular standpoint [1]. GBM stands out as the most common, aggressive, and lethal primary brain tumor in adults, known for its high vascularity and malignancy, characterised by significant neovascularization and brain parenchymal invasion [2, 3]. Despite multimodal treatments such as maximal surgical resection, radiation, and chemotherapy, glioma recurrence and death rates are significant [4, 5]. Chemotherapy is a primary cancer treatment that promotes cytotoxicity at the site of the tumour and reduces its mass [6, 7]. Conventional chemotherapeutic drugs such as doxorubicin (DOX) and temozolomide (TMZ) may be used to overcome the challenge of eradicating metastasized cancer [8], which has been found to significantly improve both the median and 5-year survival rates [5]. Researchers are interested in potential non-invasive adjuvant treatments, including drug delivery technologies using as ultrasound (US), that might improve therapeutic efficacy [9], since these chemotherapeutics alone were insufficient to meet the end objective [2, 5, 8].

TMZ is an alkylating agent employed in the treatment of GBM that promotes cell death through oxidative stress and genotoxicity [2, 10]. Due to its ability to cross the blood-brain barrier (BBB), TMZ, a first drug approved by the FDA, has been one of the most often prescribed anti-glioma drugs with limited adverse effects for the last two decades [5, 10]. However, the therapeutic benefits of TMZ are insufficient. The DNA repair protein O6-methylguanine-DNA methyltransferase (MGMT), linked to tumor resistance against alkylating drugs, is present in GBM and plays a role in conferring resistance to TMZ [11, 12]. On the other hand, due to the stem-like properties of tumour cells and genetic anomalies in GBM, which also lead to resistance to TMZ and progression [2]. Multiple trials have combined TMZ with radiation, other chemotherapeutic drugs, and phytochemicals to enhance its efficacy, however many patients continue to develop drug resistance [2]. In this research we aim to enhance the efficiency of GBM treatment by decreasing TMZ cytotoxicity resistance using US [5], and to use DOX as a reporter, to study further the sonoporation effects with a theranostic agent. US has promise as a cancer therapy since it is feasible to target specific anatomical locations with fewer side effects [8].

Ultrasound is employed in both the diagnostic and therapeutic fields. Diagnostic ultrasound is a non-invasive imaging technology that uses ultrasound waves to view inside organs. The therapeutic applications of ultrasound fall into two categories: non-destructive mechanical heating of tissue to trigger the natural physiological response to injury, as well as for selective destruction of target tissues and organs to enhance efficacy and selectivity [13]. Due to these beneficial biological interactions, US has developed a rising interest in the subject of drug delivery in recent decades [9].

US has been investigated in clinical studies for the transitory opening of the BBB, with trials to far proving the approach’s safety and viability [14–16]; it has emerged as a safe, feasible strategy for non-invasively, locally, and reversibly opening the BBB [8, 17, 18]. Researchers were also able to open the BBB using focused US with microbubbles (MBs) to enhance the delivery [18] with complex phenomena that involve the oscillation and destruction of MBs [8, 16]. Local fluid dynamics resulting from oscillating and/or collapsing cavitation bubbles are suggested to initiate the intracellular delivery mechanism induced by US [9, 19]. The plasma membrane of a cell may be torn apart by shearing or the direct contact of fluid with the cell, creating a transient pores [9, 20]. Through these pores, molecules may access the cytosol. A technique requiring temporarily increasing intracellular calcium levels is used by cells to transport intracellular vesicles to the site of damage and promptly repair these holes. Under optimal circumstances, cells survive this process, retaining a significant number of intracellular molecules and seeming to regain normal function within a few hours [9, 19]. In some situations, cellular damage is too severe, and cells die via necrosis or programmed death [19]. Researchers used the sonoporation capability of US in conjunction with several chemotherapeutics [21, 22], short hairpin RNA [22, 23], antibodies [22, 24], genes [22], and viruses [22, 25] delivery.

Increased vascularization, cell heterogeneity, self-renewing cancer stem cells, and interactions within the tumour-microenvironment (TME) all contribute to GBM progression [3]. 3D tumour spheroids provides more accurate models to explore the association between the TME, tumour reoccurrence, and drug resistance [3]. In 3D tumour spheroids, cell-cell interactions and interactions with the cell-extracellular matrix (ECM) component, enable cells to thrive *in vitro* in an environment that closely replicates *in vivo* settings [3]. Therefore, in this current study, we employed 3D tumour spheroid models to more accurately simulate and investigate the impact of US on human GBM and human epidermoid carcinoma. Typically, drug-combination assessments using US for the treatment of GBM are performed in 2D cell cultures. In this work, for the first time, the therapeutic impact of the US 96-probe device and the synergistic potential of a specific drug-combination on diffusion through a tumour, uptake by cells in a tumour sphere, and ultimately cytotoxicity comparisons in 3D GBM spheroids with 2D cultures were examined.

## 2. Materials and Methods

### 2.1. Chemicals

All chemicals utilized in this research were supplied by Sigma-Aldrich - Merck Group (Arklow, Ireland), unless stated otherwise.

### 2.2. 2D cell culture

The human GBM cell line (U-251 MG, formerly known as U-373 MG-CD14) was a gift from Michael Carty (Trinity College Dublin), and the human epidermoid carcinoma (A431), was purchased from an ATCC European distributor (LGC standards). The absence of mycoplasma is checked by using the MycoAlert PLUS Mycoplasma Detection kit (Lonza). Cells were maintained in Dulbecco’s Modified Eagle Medium (DMEM)-high glucose without sodium pyruvate supplemented with 10 % Fetal Bovine Serum (FBS) and 1 % penicillin/streptomycin. Cells were maintained in a humidified incubator containing 5 % CO_2_ atmosphere at 37 ᵒC in TC flask T25 (Sarstedt), standard for adherent cells. Cells were routinely sub-cultured when 70 - 80 % confluency was reached using a 0.25 % w/v Trypsin-EDTA solution [26].

### 2.3. 3D cell culture

U-251 MG human GBM and A431 human epidermoid carcinoma cells were used to develop 3D tumour spheroids. Single cell suspension (with desire seeding density) seeded in Nunclon™ Sphera™ 96-Well -low attachment plates (Thermo Fisher Scientific) in DMEM-high glucose supplemented with 10 % FBS and 1 % penicillin/streptomycin [27, 28]. Centrifuge the low attachment plates at 1250 rpm for 5 minutes for U-251 MG cells, 4000 rpm for 10 minutes for A431 cells, and transfer the plates to an incubator (37 °C, 5 % CO_2_, 95 % humidity). Fresh media is added every second day by replenishing old media in each well without disturbing tumour spheroids. Tumour spheroids formation was observed within 3 and 4 days for A431 and U-251 MG, respectively [28]. Tumour spheroid formation was visually confirmed daily using an Optika XDS-2 trinocular inverse microscope equipped with a Camera ISH500, and their mean diameters were analysed using “ImageJ version 1.53.e” software [26].

### 2.4. Ultrasound probe device

The ultrasonic liquid processor (Figure 1A) used in this research was the VCX 750 [Dimensions H x W x D: (235 x 190 x 340mm)]. It’s robust and adaptable, processing a wide range of sample types and volumes for a variety of applications. The CVX 750 processor’s maximum power output is 750 watt and its frequency is 20 kHz. This US device consists of a Ti6Al4V titanium alloy standard probe with a 13 mm tip diameter and a 139mm replaceable tip length. We modified this by fixing a 96 well probe at the end of the replaceable tip. This 96 well probe is designed by using Stainless steel, diameter of a tip is 2 mm and tip length is 20 mm. The 96-probe system is designed to fit perfectly into the 96 well plate (Figure 1B), which can be used to direct tumour sphere treatment. The retort stand is used to hold the US 96 probe unit vertically. A laboratory jack is used to hold the tumour sphere grown 96 well plate and to move it towards the 96 probes. A Sonics - Vibra cell power unit is needed to fix into this US 96 probe. Using the power unit, the US treatment time (0 to 99 min), temperature, pulser, and amplitude can be controlled. Experiments found that the best parameters to use are 20 % amplitude, pulses of 59 seconds on and 1 second off, with different time ranges (1, 3, 5, 10, and 20 min). During the treatment, place the sample (96 plate without lid) in the centre of the laboratory jack and slowly move it into the 96 probe. A 2 mm distance must be kept from the bottom of the well to the probe tip during the treatment. After setting up the unit, the device is turned on. The samples are treated with US using the parameters that were set. The part of the protocol described here, is published on protocols.io, dx.doi.org/10.17504/protocols.io.e6nvwkpdwvmk/v1.

**Figure 1:**
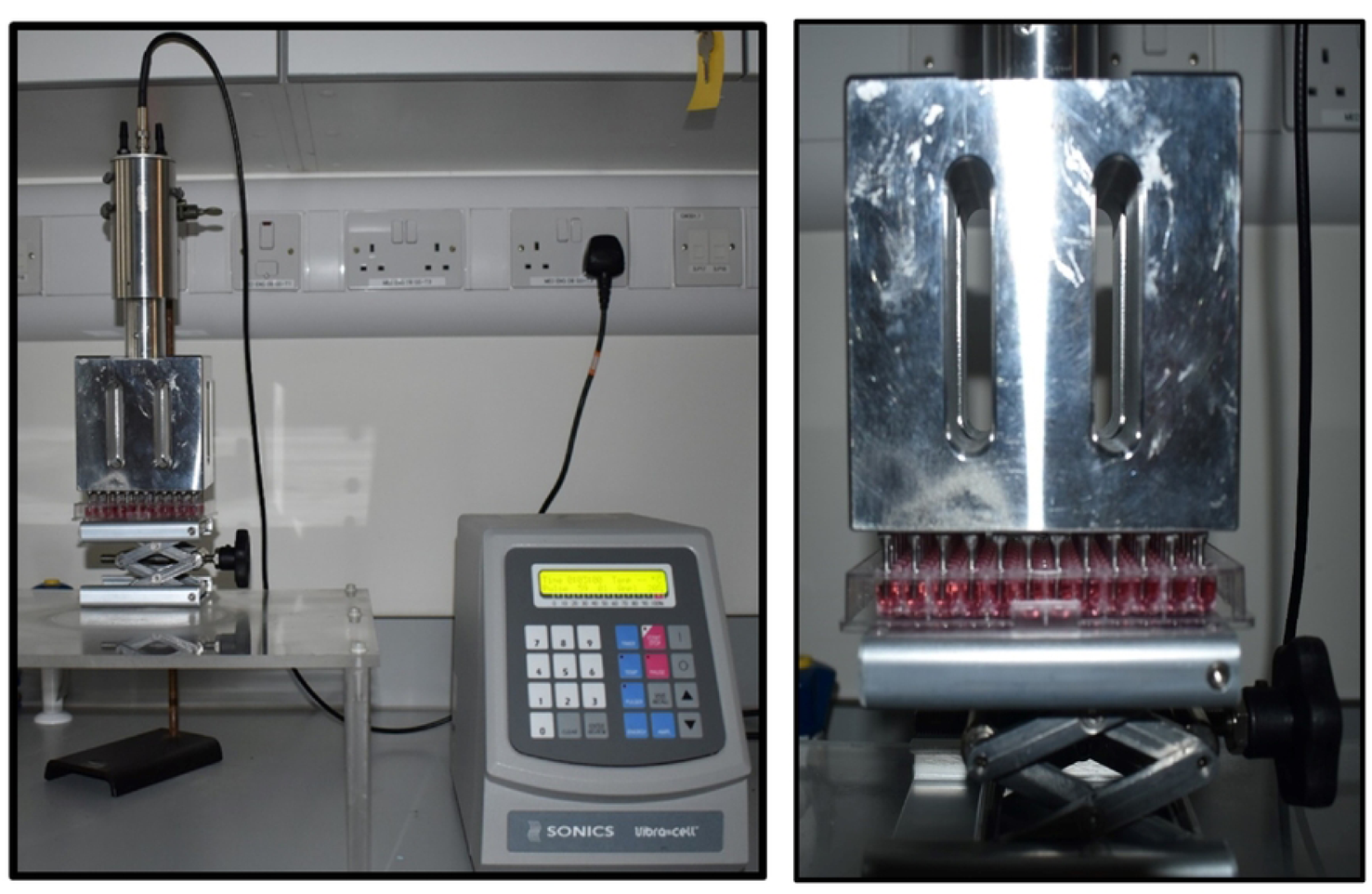
Ultrasound probe system. A) Image of US probe system B) Image of the probes demonstrating the position of the microplates for cell treatment.

### 2.5. Ultrasound treatment

U-251 MG and A431 cells were seeded at a density of 1 x 10^4^ cells/well (100 µl culture medium per well) into flat bottom 96-well plates (Sarstedt, Ltd) and were seeded at a density of 1 x 10^4^ cells/well (200 µl culture medium per well) into Nunclon™ Sphera™ 96-well -low attachment plates (Thermo Fisher Scientific) respectively, for 2D and 3D cell culture. Cells were incubated overnight at 37 ᵒC in a humidified atmosphere. DMEM media without sodium pyruvate supplemented with 10 % FBS and 1 % penicillin/streptomycin were used throughout this study. In 2D cell culture, after 24 h of incubation, checked 70 - 80 % confluency, 50 µL of media were removed, leaving 50 µL of media for treatment in each well.

In tumour sphere culture, after 3 days of incubation for A431 and 4 days of incubation for U-251 MG, 100 µL of media were removed, leaving 100 µL of media for treatment in each well, unless otherwise specified. Cells/tumour spheres were then treated with an US probe system at six different time points (0, 1, 3, 5, 10, and 20 minutes) using a Sonics - Vibra cell power unit, fixed into an US 96 probe. Using the power unit, the US treatment time (0 to 99 min), temperature, pulser, and amplitude can be controlled.

Experiments found that the best parameters to use are 20 % amplitude, pulses of 59 seconds on and 1 second off, with different time ranges (1, 3, 5, 10, and 20 min). 50 µL (for 2D) and 100 µL (for 3D) of fresh culture media were added immediately following the US treatment, and cells were incubated at 37 ℃ using 5 % CO_2_ for 24, 48, 72, and 96 hours. Dimethyl sulfoxide (DMSO) (20 %) was used as a positive control. Multiple US treatments (single, double, and triple) were carried out for tumour spheres as where it is mentioned. The multiple US treatments are a combination of individual US treatments with a 24 hours incubation gap between each treatment.

### 2.6. TMZ and DOX treatment

Stock solutions of compounds were dissolved in DMSO and stored at -20 °C. These stocks were subsequently used to make the working solutions in media. The highest concentration of DMSO used was 0.5 %. For 2D cell culture, cells were seeded at a density of 1 × 10^4^ cells (48 hours) and 2.5 × 10^3^ (144 hours) exposure time period with 100 µL media per well in 96-well plates (Sarstedt). For 3D cell culture, cells were seeded at a density of 1×10^4^ cells/well (200 µl of culture medium per well) into Nunclon™ Sphera™ 96-well low attachment plates (Thermo Fisher Scientific).

Plates were incubated at 37 °C with 5 % CO_2_ to allow proper adherence (2D) and tumour sphere formation (3D) as explained above. Existing media were removed from each well and cells / tumour spheres were treated with TMZ/DOX at varying concentrations (500 – 0.976 µM; 100 µL/well for 2D cells ; 200 µL/well for 3D cells), 20 % (v/v) DMSO was used as a positive control, and 0.5 % (v/v) DMSO as a negative control, and incubated at the appropriate time point. No deleterious effects were observed from the negative control solvent. Another set of plates were treated with US using the same procedure as in section 2.5.

### 2.7. Cell viability assay

Cell viability was assessed using the Alamar Blue™ cell viability reagent (Thermo Fisher Scientific). Following exposure at specific time points (48 or 144 hours), cells were rinsed once with sterile phosphate-buffered saline (PBS) (Merck, Ireland). Subsequently, a 10% Alamar Blue™ solution in DMEM (without FBS and antibiotics) was added to each well and incubated at 37 °C for 3 hours. Fluorescence measurements (excitation at 530 nm; emission at 590 nm) were performed using a Varioskan™ LUX multimode microplate reader (Thermo Fisher Scientific). All experiments were performed at least three independent times with a minimum of three replicates per experiment and are presented as mean ± S.E.M. The data (in fluorescence units from the microplate reader) for the test wells were normalised to the assay control, and cell growth was calculated as a change in viability over time [26].

### 2.8. Cell viability assay (3D cell culture)

Cell viability was analysed using the Alamar Blue™ cell viability reagent (Thermo Fisher Scientific). After the post treatment incubation tumour spheres were washed with sterile phosphate buffered saline (PBS), trypsinization using 0.25 % w/v Trypsin-EDTA solution and incubated for 3 hours at 37 °C with a 10 % Alamar Blue™ solution. Fluorescence was detected using an excitation wavelength of 530 nm and an emission wavelength of 590 nm on a Varioskan Lux multiplate reader (Thermo Scientific). All experiments were performed at least three independent times with a minimum of 30 replicates per experiment [26].

### 2.9. Live/Dead cell staining using propidium iodide (PI)

Tumour spheres were constructed using Nunclon™ Sphera™ 96-well -low attachment plates (Thermo Fisher Scientific) and US treated using the US probe system as previously described here. After US treatment immediately replenished media in wells and incubated were incubated at 37 ℃ using 5 % CO_2_ for 24, 48, 72 and 96 hours. For PI staining the media was removed and the tumour spheres washed with PBS and trypsinized using 0.25 % w/v Trypsin-EDTA solution into single cell suspension. Then inactivate trypsin and collect cells from each treatment point into single centrifuge tube for centrifugation at 250 x g for 5 minutes. The supernatant was then aspirated, and the pellet re-suspended in 1 ml of 1X PBS. PI was then added to the cell suspension at 10 µg/mL and incubated for 1 minute in the dark. PI fluorescence was then measured using a Beckman Coulter CytoFLEX flow cytometer with a 488 nm blue laser for excitation and an FL1 standard filter for PI fluorescence, where PI binds to nuclear degradation from dead cells [26].

### 2.10. DOX sonoporation analysis

Doxorubicin hydrochloride (DOX.H) (Fisher Scientific) is a chemotherapeutic agent with intrinsic fluorescence, with excitation/emission of 470 / 585 nm. Stock solutions (1mg/5ml) of compounds were dissolved in DMSO and stored at -20 °C. A431 and U-251 MG tumour spheres were seeded in a Nunclon Sphera 60mm dish (Thermo Fisher Scientific) at a density of 1 × 10^5^ cells/ ml using DMEM high glucose in the absence of sodium pyruvate and incubated at 37 °C using 5 % CO_2_ for 3 to 4 days, respectively. Subsequently, the culture medium was removed and the tumour spheres were washed with 1× PBS. Tumour spheres were replenished with DMEM media, without sodium pyruvate and phenol red, containing 10 µg/ml DOX.H. Subsequently dishes with tumour spheres were treated with US for 1 and 3 minutes, as described in the section 2.5. Then all the dishes (tumour spheres without US and DOX, with only US, with DOX and without US, DOX with 1 min of US, DOX with 3 min of US) were incubated for 24 h at 37 °C using 5 % CO2. After the post treatment incubation, tumour spheres were trypsinized into a single cell suspension and centrifuged at 1200 rpm for 5 min. The cells were suspended in 1× PBS, and their fluorescence was analyzed using a Beckman Coulter CytoFLEX Flow Cytometer. The system utilized a 488 nm blue laser for excitation and the FL1 standard filter for FITC measurement [26].

### 2.11. Statistical analysis

All experiments were conducted at least three independent times. Curve fitting and statistical analysis were performed using Prism 6.01, GraphPad Software, Inc. (USA). Non-linear regression was employed to measure the dose-response curve. Data are presented as the % and error bars of all figures are presented using the standard error of the mean (S.E.M), multiple comparison analysis were performed using Sidak’s test, unless otherwise stated. Flow cytometry analysis was conducted using CytExpert software, with FITC-A mean values used to create statistical columns. PI uptake studies were analysed using a Two Way ANOVA with Tukey’s post-test [26].

## 3. Results and discussion

### 3.1. US probe presents cytotoxicity towards GBM and Epidermoid carcinoma cells in a time and dosage dependent manner

The TME plays a significant role in shaping tumor development, metastasis, angiogenesis, resistance to cytotoxicity, and modulation of immune cells [3, 26, 29]. In 3D cell culture, cells engage in physiological interactions, including cell-cell and cell-ECM interactions. These enable cells to thrive *in vitro*, replicating the TME found in *in vivo* GBM conditions [3, 26, 30]. The low attachment plate method was used in this study for the development of *in vitro* U-251 MG and A431 tumour spheroids and we successfully generated consistent tumor spheres that closely emulate the natural *in vivo* environment, encompassing shape and cellular responses [30, 31].

We employed a 3D cell culture model to investigate the dispersion of cytotoxic reactive species and chemotherapeutics within the tumor sphere, assess the rate of cell death, and examine the impact of both single and multiple US treatments on cell-cell and cell-ECM interactions [26]. The effects of US on cancer cells have primarily been studied using 2D monolayer cell cultures [9, 20], and although an increasing number of studies are now utilizing animal models [21, 24]. We believe that this is the first time that the US 96 probe approach has been reported for drug diffusion through a tumour sphere, uptake by cells in a tumour sphere, and ultimately induced cytotoxicity in 3D tumour spheroids compared to 2D monolayer cells.

In this study, we initially compared the impact of the US probe on both U-251 MG human GBM and A431 human epidermoid carcinoma cells, in both 2D and 3D cultures. This evaluation was conducted in DMEM high glucose supplemented with 10% FBS medium, following 24 hours of incubation. For U-251 MG 2D cells, an IC_50_ of 162.9 min (129.5 ± 205.0 min), 115.2 min (90.19 ± 147.3 min) and 22.78 min (22.19 ± 23.39 min) were found, while an IC_50_ of 26.48 min (25.35 ± 27.66 min), 21.80 min (20.95 ± 22.69 min) and 12.04 min (11.82 ± 12.27 min) were determined for U-251 MG 3D tumour spheres during single, double and triple US treatments respectively (Figure 2A).

**Figure 2:**
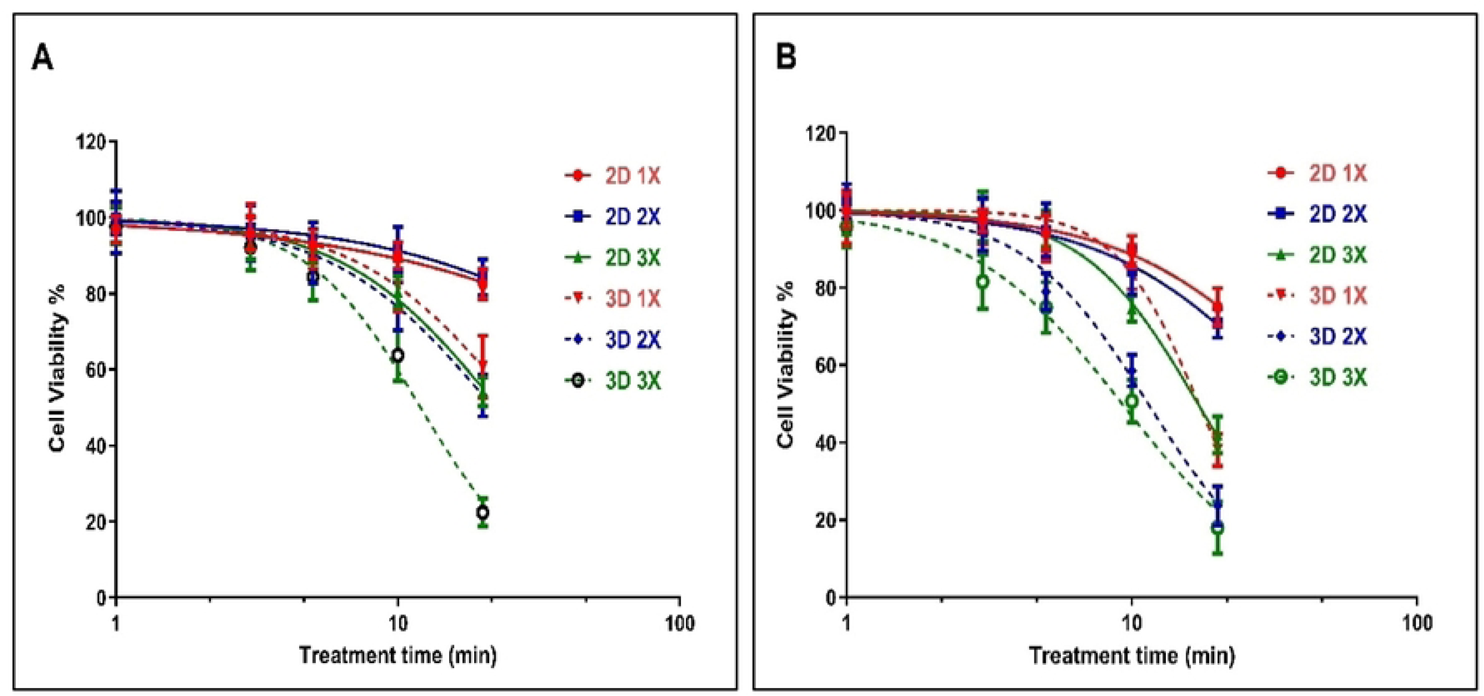
GBM cell line U-251 MG and epidermoid carcinoma cell line A431 US probe treatment. A) Comparison of 2D and 3D U-251 MG US treatments (single, double and triple) and post treatment incubation at 24 h B) Comparison of 2D and 3D A431 cell cytotoxicity induced by US (single, double and triple) and post treatment incubation at 24 h

During studies of A431 human epidermoid carcinoma, an IC_50_ of 46.0 min (42.46 ±49.83 min), 38.05 min (35.07 ± 41.29 min), and 17.05 min (16.77 ± 17.33 min) were found for A431 2D cells while an IC_50_ of 17.14 min (16.87 ± 17.43 min), 11.40 min (11.20 ± 11.60 min), and 9.239 min (9.018 ± 9.466 min) were found for A431 3D tumour spheres during single, double, and triple US treatments, respectively (Figure 2B). The results from the Two-way ANOVA indicate a significant difference in cell viability for both U-251 MG and A431 cells, both in 2D and 3D cultures, concerning varying doses of US exposure (p <0.0001). Further details regarding Tukey’s multiple comparisons test can be found in the supporting information section.

U-251 MG and A431 3D tumor spheres exhibited lower resistance to US therapies compared to their 2D cells across single, double, and triple treatments. This US treatment sensitivity in 3D tumour spheres might be influenced by the differences in cellular organization, the added dimension, polarity, and geometry inherent in 3D spheroids [3, 32]. US treatments were also able to effectively induce 2D cell and 3D tumour sphere cell death in a dosage and frequency of treatment dependent manner in both U-251 MG and A431. Nevertheless, U-251 MG cells exhibited greater resistance to the treatment in comparison to the A431 cell lines.

Following that, we assessed the cytotoxic effects of both single and multiple US treatments on GBM tumor spheres at various incubation periods. An IC_50_ of 75.22 min (58.16 ± 97.29 min), and 26.41 min (25.55 ± 27.31 min) were found for U-251 MG tumour spheres during single and fivefold US treatments, respectively, with 8 h post treatment incubation (Figure 3A). An IC_50_ of 24.87 min (24.35 ± 25.39 min), 9.18 min (9.08 ± 9.29 min), (Figure 3B) and 30.95 min (29.55 ± 32.42 min), 7.91 min (7.78 ± 8.04 min), (Figure 3C) and 57.91 min (44.60 ± 75.20 min), 8.46 min (8.33 ± 8.60 min) (Figure 3D) were found for U-251 MG tumour spheres during single and fivefold US treatments with 24 h, 48 h and 72 h post treatment incubations, respectively. Finally, an IC_50_ of 42.73 min (38.81 ± 47.05 min), 7.96 min (7.85 ± 8.08 min), (Figure 3E) and 15.48 min (15.10 ± 15.87 min), 6.00 min (5.88 ± 6.12 min) (Figure 3F) were found for U-251 MG tumour spheres during single and fivefold US treatments with 96 h and 120 h post treatment incubations, respectively. A full description of the IC_50_ values and ranges during 1X, 2X, 3X, 4X, and 5X US treatments are shown in the supplementary section as a Table S1.

**Figure 3:**
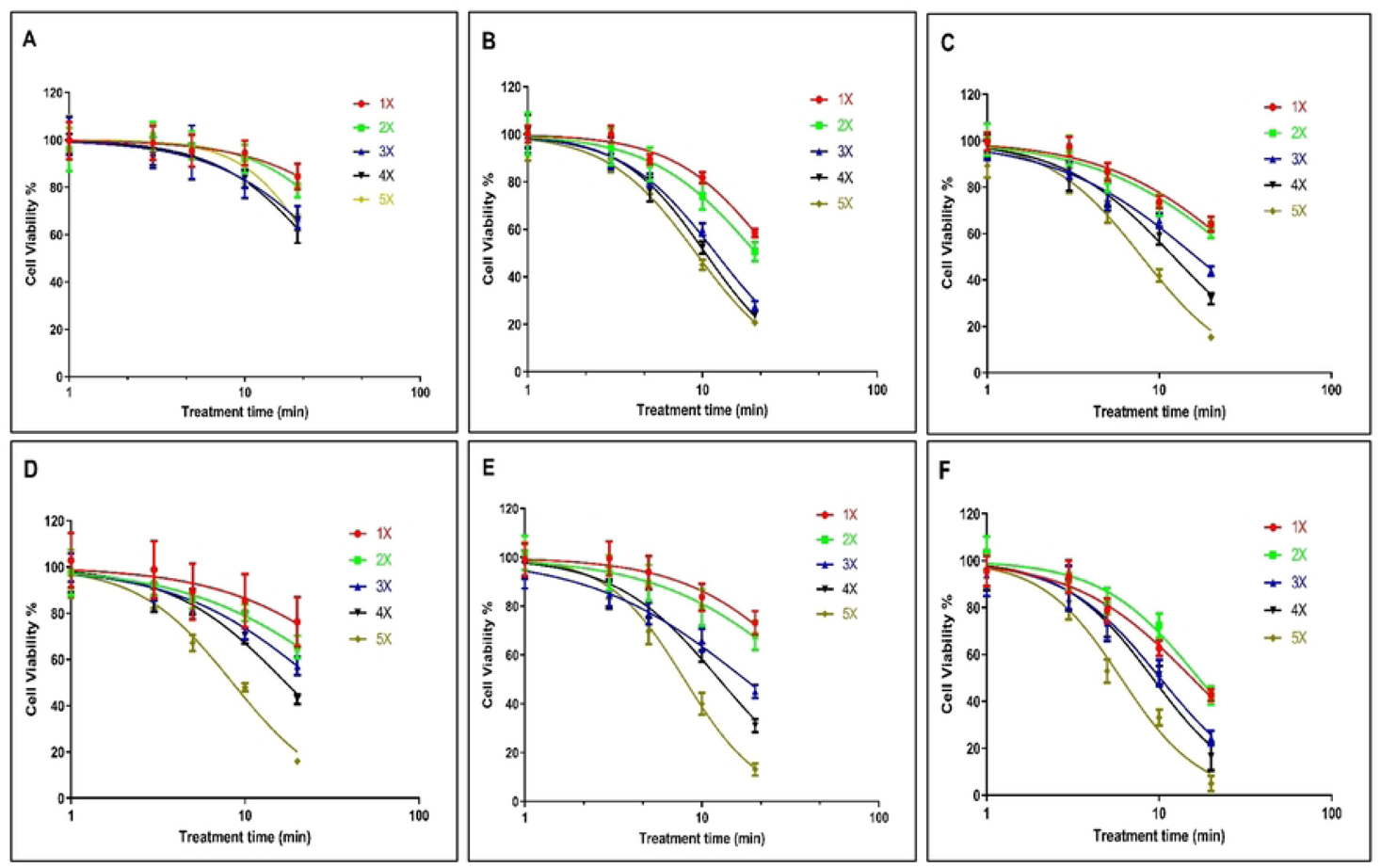
U-251 MG tumour sphere single and multiple US treatments with different incubations. (A) 8 h, (B) 24 h, (C) 48 h, (D) 72 h, (E) 96 h and (F) 120 h post treatment incubations.

A two-way ANOVA revealed a significant disparity in viability concerning the doses of single and multiple US treatments, as well as different post-treatment incubation periods (p < 0.0001). Additional details regarding Tukey’s multiple comparison test can be found in the supporting information section.

Based on the cytotoxicity findings illustrated in Figure 3, it is evident that even with the longest exposure time (20 min) and extended incubation period (120 h), a single US exposure did not lead to complete cytotoxicity in the tumor spheres. According to Figure 3F, the cell viability remained at 42.56%. Given that a single US treatment failed to achieve significant cytotoxicity and prevent tumor regrowth, we hypothesised that employing multiple (up to five consecutive daily) US treatments would result in more favourable outcomes. The hypothesis proved according to the data, the lowest cytotoxicity observed during single US treatment with 8 h post treatment incubations, while highest cytotoxicity observed during five consecutive daily US treatments with 120 h post treatment incubations. These findings affirm that multiple US treatments substantially triggered cytotoxicity within the tumor spheres. In the case of U-251 MG, these treatments either fully or partially reversed the regrowth ability of the tumour spheres. These results bear significant implications for upcoming animal model studies and human trials, suggesting that single US treatments might prove inadequate in delivering substantial benefits.

According to Paškevičiūtė et al., 2020, US has no impact on the viability of MDA-MB-231 triple-negative breast cancer cells or A549 non-small cell lung cancer cells [9]. In line with these results, we did not detect any significant cytotoxicity effect at the single US treatment up to 2 minutes of US exposure.

Our results demonstrate that US alone is capable of inducing cytotoxicity when cells/tumour spheres are exposed to prolonged durations of US, and that multiple treatments augment these effects. US-treated tumour spheres incubated for an extended duration (120 h) exhibited a notable reduction in cell viability and significantly more cell death compared with 8 h. When US exposure length and treatment frequency are increased, persistent pore formation in membranes, lysis, and ultimately cell death and tumour sphere damage result [9, 20]. The response to the US treatment exhibited a distinct kinetic pattern over time, differing significantly from the response observed with cold atmospheric plasma (CAP) treatment [26]. In our previous study, we used U-251 MG tumour spheres and CAP treated to evaluate the cytotoxicity response. Interestingly, we observed considerably higher cell death in short-term post treatment incubations compared with long-term incubations [26]. Ultimately, the analysis demonstrates that the US 96-probe device can induce cytotoxicity in tumour spheres in a manner that depends on both the dosage and duration of exposure.

#### Impact of US treatment on cell membrane damage within tumour spheres

PI staining was employed to confirm the cell death and cytotoxicity induced by the US 96-probe in U-251 MG tumour spheres. PI is a fluorescent, nucleic acid intercalating agent that cannot penetrate intact cell membranes, serves to identify dead cells with compromised plasma membranes within a tumour sphere population [26]. PI uptake was assessed 24 h after single (Figure 4A), double (Figure 4B), and five consecutive daily (Figure 4C) US treatments. The percentage of cells permeable to PI increased to almost 40 %, 45 % and 80 %, respectively, following single, double, and five multiple US treatments for 20 minutes, as shown in Figure 4D. This also demonstrates that US treatment can damage the tumour sphere’s cell membrane and induce cytotoxicity. This validates the results obtained from the Alamar Blue™ cell viability assay. Two-way ANOVA showed a significant variance in PI uptake between 5 and 10 min of US doses for both double and five multiple US treatments (p < 0.0001). Additionally, a significant difference was observed at 20 min for both single and multiple US treatments (Figure 4D). A full description regarding Tukey’s multiple comparisons test can be found in the supporting information. Validating our findings, researchers discovered that US inhibited spheroid growth, metabolic activity, disrupted spheroid integrity, and increased DNA double strand breaks, leading to damage in human prostate cancer (PC-3) and GBM (U87) cell lines [33].

**Figure 4:**
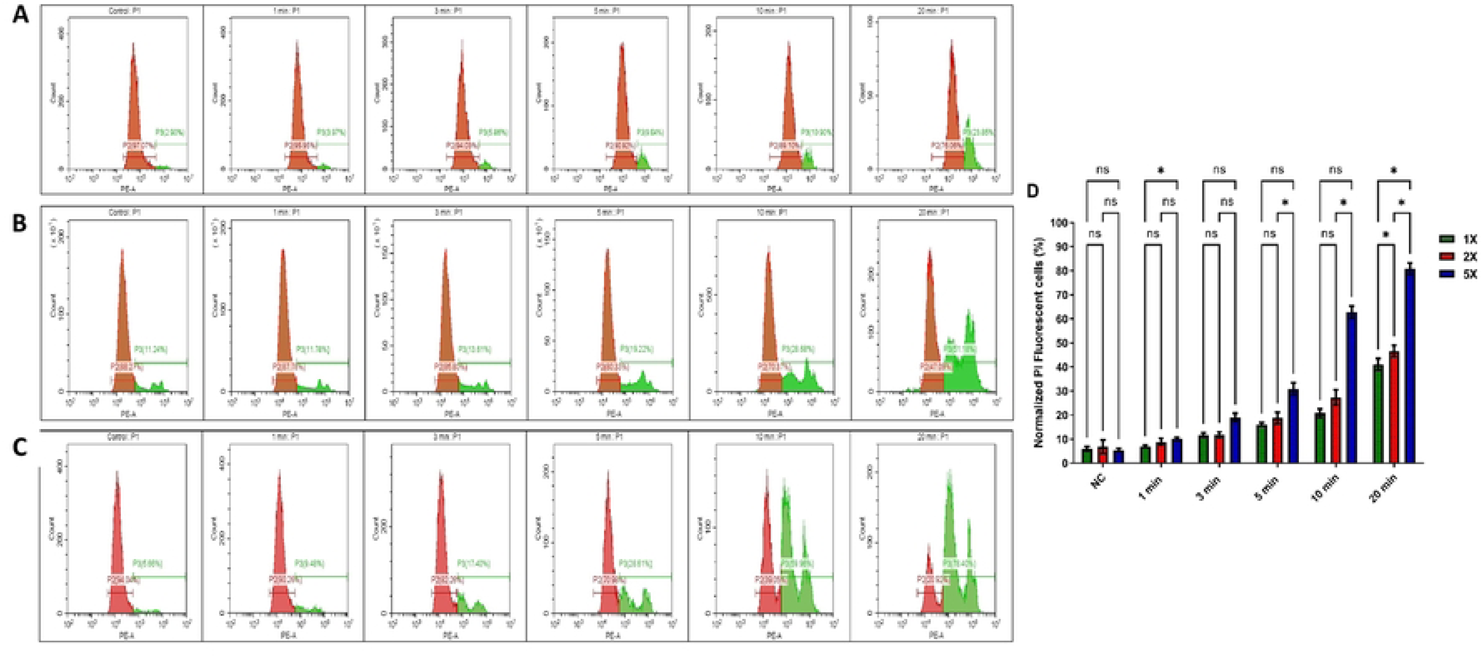
PI uptake in ultrasound probe treated U-251 MG tumour spheres. PI uptake was quantified through flow cytometry and served as an indicator of cell death. Cells were treated for 1, 3, 5, 10, and 20 min. PI uptake was subsequently measured 24 h post treatment in both (a) single US treatment, (b) Double (2X) US treatments, (C) 5X US treatment, (Positive control can be seen in the supporting information as Figure S1.) (d) The normalized PI uptake was then assessed 24 h after single and multiple treatments and depicted in a bar chart. The panel A-C are representative of 3 replicates. All data points showed statistical significance, with the exception of the control and 1 min treatment times. ns, not significant, *p ≤ 0.05; ****p ≤ 0.0001

#### Tumour spheres morphological changes induced by ultrasound probe

The alterations in the morphology of tumour spheres caused by US treatments were investigated to enhance our comprehension of the cellular death mechanisms involved. After the initial US treatment, the diameter of U-251 MG tumour spheres showed a notable reduction for both 10 and 20 minute treatment durations (Figure 5B). Representative images of tumour sphere images depicting the morphological transformations caused by 20 min of US treatment for U-251 MG as shown in Figure 5A. Repetitive ultrasound treatments led to notable cumulative cytotoxic effects, evident through spheroid rupture, shrinkage, and significantly impaired tumor regrowth capability. These effects were more pronounced with extended overall US treatment duration. Consequently, it appears that for clinical applications, a series of US treatments over a comparatively extended timeframe would likely be essential. Interestingly, tumour spheres reduced in size after the second US treatment and gradually broke apart on consecutive treatments. Following the initial US treatment and 24 h of incubation, there were no notable morphological changes observed in the U-251 MG tumour spheres (Figure 5A-II), in comparison to their pre-treatment state (Figure 5A-I). Then tumour spheres swelled, ruptured significantly after the second (Figure 5A-III) and third (Figure 5A-IV) treatments, and disintegrated after the fourth (Figure 5A-V) and fifth (Figure 5A-VI) treatments.

**Figure 5:**
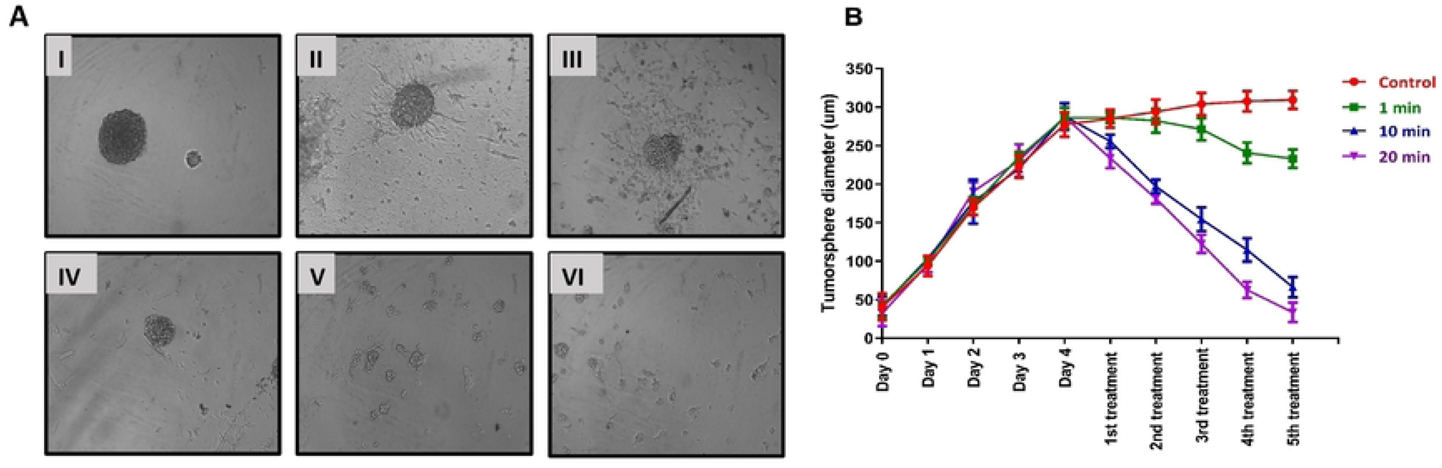
U-251 MG tumour sphere morphology and size (diameter) variation analysis followed by US treatment. A) U-251 MG tumour sphere morphological changes with 20 min US treatment (I-Before US treatment, II – 24 h after 1st treatment, III – 24 h after 2^nd^ treatment, IV – 24 h after 3^rd^ treatment, V – 24 h after 4^th^ treatment, VI – 24 h after 5^th^ treatment) B) U-251 MG tumour sphere size (diameter) followed by different US treatments.

Overall, the results were consistent, demonstrating increased cytotoxicity and the inability to reform tumour spheres after multiple US treatments. Multiple US treatments might potentially disrupt cell–cell and cell–ECM interactions. This disruption could lead to a reduction in the volume of densely packed tumor spheres, consequently causing a decrease in their diameter. A two-way ANOVA analysis revealed a significant variance in the tumour sphere diameter across different US treatment durations (p < 0.0001). Detailed information regarding Tukey’s multiple comparison test can be found in the supporting information section.

#### Effect of US on TMZ delivery in human glioblastoma and epidermoid carcinoma cell models

Initially, we explored the effects of the 3 min US exposure on TMZ delivery into U-251 MG human GBM and A431 human epidermoid carcinoma after 144 h of post-treatment incubation time. An IC_50_ of 133.0 µM (124.8 ± 141.8 µM) and 18.16 µM (17.34 ± 19.01 µM) were found for U-251 MG 3D spheroids during TMZ treatment without and with US, respectively (Figure 6A), while an IC_50_ of 70.63 µM (65.98 ± 75.60 µM) and 13.55 µM (13.14 ± 13.98 µM) were found for A431 3D spheroids during TMZ treatment without and with 3 min US exposure respectively, after 144 h post treatment incubation (Figure 6A).

**Figure 6:**
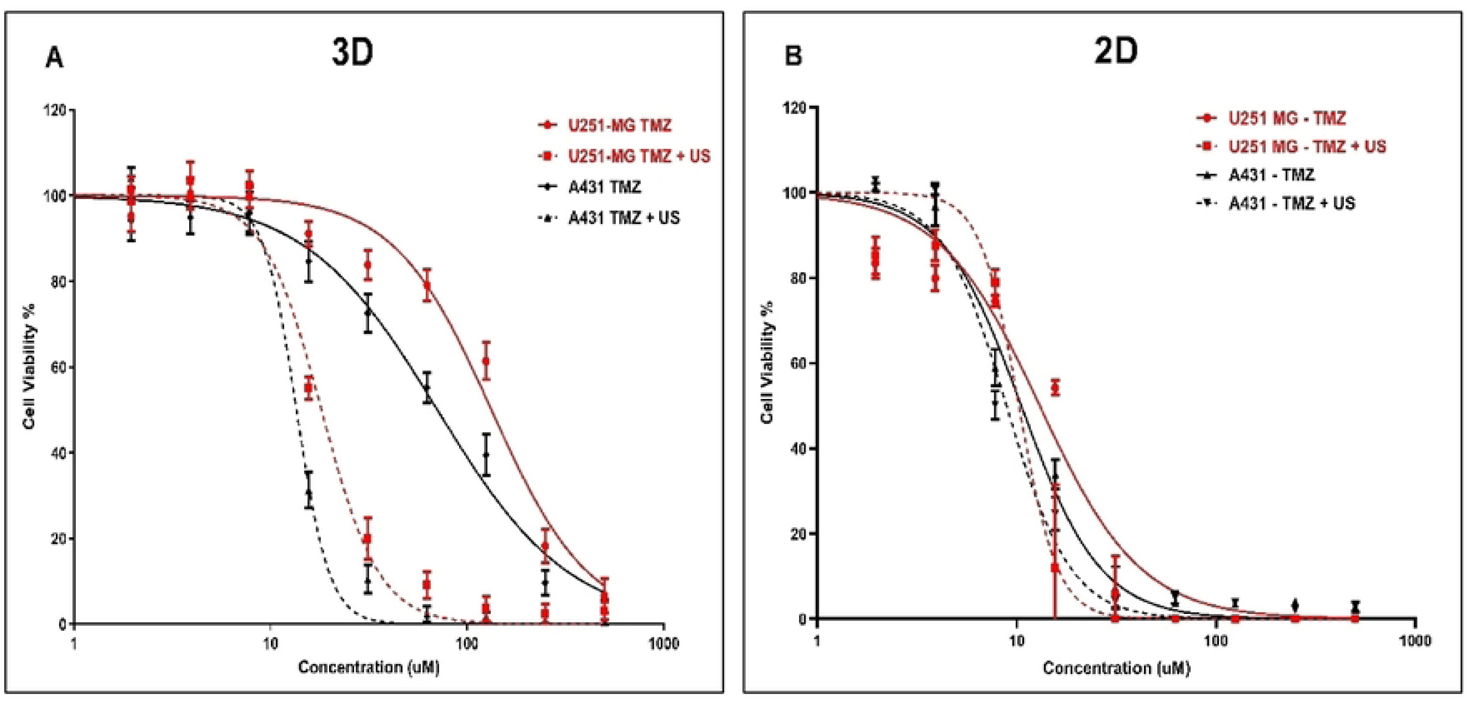
TMZ cytotoxicity analysis with and without US (3 min) combination using U-251 MG and A431 2D cells and 3D tumour spheroids A) TMZ cytotoxicity in 3D cell cultures after 144 h incubation B) TMZ cytotoxicity in 2D cell cultures after 144 h incubation.

After investigating the synergistic effect of US and TMZ on 3D tumour spheres, we then looked at its effect in U-251 MG and A431 on 2D cells with 144 h post treatment incubations. An IC_50_ of 13.06 µM (10.85 ± 15.52 µM) and 10.27 µM (9.086 ± 11.66 µM) were determined for U-251 MG 2D cells during TMZ treatment without and with US, respectively (Figure 6B), while an IC_50_ of 10.55 µM (9.728 ± 11.45 µM) and 8.898 µM (8.121 ± 9.780 µM) were found for A431 2D cells during TMZ treatment without and with 3 min US exposure respectively (Figure 6B).

Then we reduced the post treatment incubation time to 48 h to investigate the cytotoxic effects at shorter incubations (outcomes can be seen in supplementary section as a figure S2). An IC_50_ of 300.1 µM (272.1 ± 331.1 µM) and 201.8 µM (185.4 ± 219.7 µM) were determined for U-251 MG 2D cells during TMZ treatment without and with US, respectively (Figure S2A), while an IC_50_ of 180.9 µM (172.8 ± 189.3 µM) and 98.53 µM (93.59 ± 103.7 µM) were found for A431 2D cells during TMZ treatment without and with US, respectively (Figure S2A). Interestingly, an IC_50_ of 707.6 µM (629.5 ± 795.3 µM) and 264.9 µM (256.7 ± 273.4 µM) were determined for U-251 MG 3D spheroids during TMZ treatment without and with US, respectively (Figure S2B), while an IC_50_ of 288.7 µM (274.4 ± 303.7 µM) and 195.2 µM (186.3 ± 204.5 µM) were found for A431 3D spheroids during TMZ treatment without and with 3 min US exposure, respectively after 48 h post treatment incubation (Figure S2B).

It is noteworthy that the sensitivity of TMZ was significantly enhanced after the US treatment in the U-251 MG and A431 on 3D tumour spheroids relative to 2D cells. When tumour spheroids were incubated with TMZ at the highest concentration (500 µM) for 48 h after US treatment, cell viability decreased approximately from 59.58 % to 14.31 % in U-251 MG; and from 30.15 % to 9.37 % in A431 tumour spheres (Figure S2B). This pattern of TMZ cytotoxicity intensified after 144 h of incubation. At doses of TMZ greater than 15 µM, where a four-fold increase in cytotoxicity can be observed in tumour spheres, while only around one-fold increase in cytotoxicity can be observed in 2D cells. When tumour spheroids are incubated with TMZ 62 µM for 144 h after US treatment, cell viability decreases approximately from 79.11 % to 9.21 % in U-251 MG and from 55.25 % to 2.35 % in A431 tumour spheres (Figure 6A).

It was also found that there was an increase in TMZ cytotoxicity in longer post-treatment incubations compared to shorter incubations. This evidence demonstrates the ability of US to increase cytotoxicity with TMZ. The intracellular redox status and the decrease and activation of antioxidant-related signalling molecules determine TMZ resistance. Combining TMZ with US seems to minimise this TMZ resistance and may provide a solution to TMZ cytotoxicity and resistance. [2, 34]. This results also demonstrates the important of adopting 3D cell culture models in pre-clinical research to get accurate outcome from toxicological assessments. Overall, our findings suggest that the combination of US and TMZ represents a promising approach to enhance the effectiveness of chemotherapy for both GBM and epidermoid carcinoma.

#### Effect of US on DOX delivery in human glioblastoma and epidermoid carcinoma cell models

To elucidate the mechanism further, the theranostic chemotherapeutic agent DOX is used to correlate cytotoxicity with uptake into tumour cells and distribution throughout the tumour sphere. We explore the effects of the 3 min US exposure on DOX delivery into U-251 MG human GBM and A431 human epidermoid carcinoma tumour spheres after 48 h of post-treatment incubation time. An IC_50_ of 44.00 µM (41.80 ± 46.31 µM) and 4.835 µM (4.503 ± 5.190 µM) were found for U-251 MG 3D spheroids during DOX treatment without and with US, respectively (Figure 7A), while an IC_50_ of 27.18 µM (26.10 ± 28.31 µM) and 3.045 µM (2.874 ± 3.227 µM) were found for A431 3D spheroids during DOX treatment without and with 3 min US exposure, respectively, after 144 h post treatment incubation (Figure 7A).

**Figure 7:**
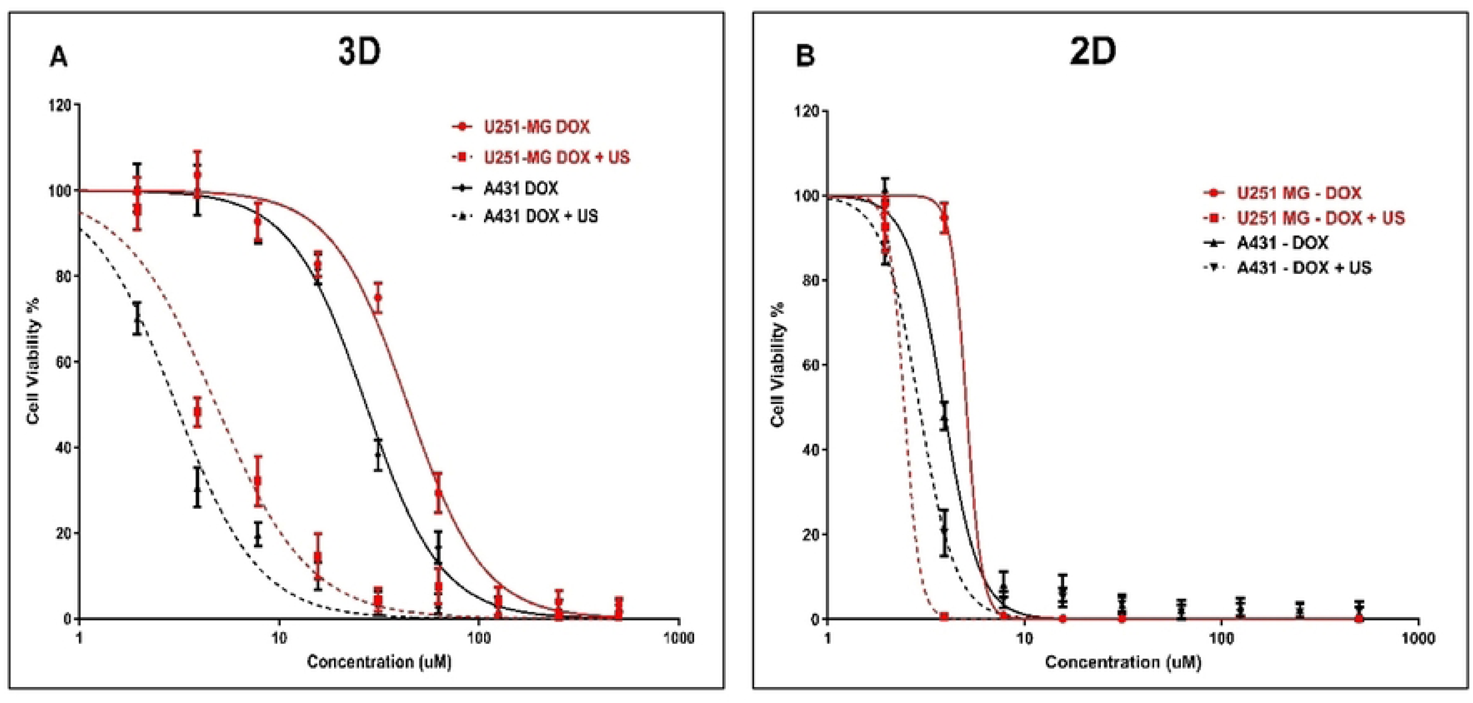
DOX cytotoxicity analysis with and without US (3 min) combination using U-251 MG and A431 2D cells and 3D tumour spheres. A) DOX cytotoxicity in U-251 MG and A431 3D cells with 144 h incubation B) DOX cytotoxicity in 2D cells with 144 h incubation.

After investigating the synergistic effect of US and DOX on 3D tumour spheres, we then extended our study to compare the activity of U-251 MG and A431 in 2D cells with 144 h post treatment incubations. An IC_50_ of 5.072 µM (4.701 ± 5.472 µM) and 2.454 µM (1.965 ± 3.065 µM) were found for U-251 MG 2D cells during DOX treatment without and with US, respectively (Figure 7B), while an IC_50_ of 3.884 µM (3.721 ± 4.055 µM) and 2.922 µM (2.786 ± 3.065 µM) were determined for A431 2D cells during DOX treatment without and with 3 min US exposure, respectively (Figure 7B).

Then we reduced the post treatment incubation time to 48 h to investigate the cytotoxic effects at shorter incubations (outcomes can be seen in supplementary section as a figure S3). An IC_50_ of 36.02 µM (27.89 ± 45.30 µM) and 25.33 µM (22.83 ± 27.96 µM) were determined for U-251 MG 2D cells during DOX treatment without US and with US, respectively (Figure S3A), while an IC_50_ of 25.73 µM (23.45 ± 28.31 µM) and 21.45 µM (20.06 ± 22.94 µM) were found for A431 2D cells during TMZ treatment without US and with US, respectively (Figure S3A). An IC_50_ of 159.4 µM (145.0 ± 175.2 µM) and 37.13 µM (35.03 ± 39.35 µM) were found for U-251 MG 3D spheroids during DOX treatment without and with US, respectively (Figure S3B), while an IC_50_ of 93.10 µM (86.89 ± 99.76 µM) and 24.08 µM (22.79 ± 25.45µM) were found for A431 3D spheroids during DOX treatment without and with 3 min US exposure, respectively, after 48 h post treatment incubation (Figure S3B).

Increasing the incubation time is the most crucial way to differentiate between the effectiveness of DOX with and without US. In some studies, it was determined that US did not improve DOX (concentration 10 µM) cytotoxicity in monolayer-cultured cancer cells and 3D tumour spheroid models with up to a 2 min US exposure and shorter post-treatment incubations, such as 1 h and 2 h [9]. Similarly, in our work, we observed no significant cytotoxicity up to 10 µM DOX with or without US for U-251 MG and A431 cells / tumour spheroids after 48 h of post-treatment incubation (Figures S3A and S3B). However, when the DOX concentration was increased above 10 µM, cytotoxicity was enhanced, and the US was able to augment DOX toxicity. We were able to induce cytotoxicity even at a lower DOX concentration, once we incubated cells/tumour spheroids for a longer period of time (144 h). When tumour spheroids are incubated with DOX 10 µM for 144 h after US treatment, cell viability approximately decreased from 90 % to 20 % in U-251 MG and from 90 % to 10 % in A431 tumour spheres.

According to our study, DOX cytotoxicity rises marginally when combined with US treatment in U-251 MG GBM and A431 epidermoid cancer in 2D cells, while showing considerably increased DOX sensitivity in 3D tumour spheroids. When combined with US, the IC_50_ in 3D GBM spheroids was reduced by more than four-fold; however, the reduction in 2D cells was a little bit above one-fold during 48 h of post-treatment incubation. This same pattern of DOX cytotoxicity was intensified over a 144 h of incubation. When combined with US, the IC_50_ in 3D GBM spheroids was reduced by more than nine-fold; however, the reduction in 2D cells was only two-fold.

This study also suggests that US may enhance the cytotoxicity of DOX. In U-251 MG and A431 3D tumour spheroids, the sensitivity of DOX was considerably increased after US therapy. When tumour spheroids are incubated with DOX at higher concentration (500 μM) for 48 h after US treatment, cell viability decreased approximately from 30.15% to 1.25% in U-251 MG, and from 13.51% to 2.39% in A431 tumour spheres. This same pattern of DOX cytotoxicity was observed over a 6 day incubation period. At doses of DOX greater than 125 μM, U-251 MG cytotoxicity increased by more than twentyfold; while A431 cytotoxicity increased by more than ten-fold, when exposed to US. When tumour spheroids are incubated with DOX 500 μM for 6 days after US treatment, cell viability approximately decreased from 30.15% to 1.25% in U-251 MG and from 13.51% to 1.13% in A431 tumour spheres.

DOX is among the most extensively utilized chemotherapeutic drugs used to treat a wide variety of neoplasms in humans [35]. It acts as a sequence-selective DNA intercalating agent, specifically targeting topoisomerase II, resulting in DNA damage and the production of reactive oxygen species (ROS), which are principally accountable for its cytotoxic effects [36]. DOX as a drug is cell cycle non-specific; however, a maximal impact has been recorded during the G0/G1 phase of the cell cycle, resulting in damage repair, cell cycle arrest, or apoptosis depending on the level of DNA damage [36, 37]. DOX has lower response rates, lower selectivity, and higher complication rates. Consequently, it is essential to optimise drug delivery to a specific cancer site while reducing the dosage for normal cells and tissues [36]. Overall, our results indicate that the combination of US and DOX is a potential combination therapy for enhancing the efficacy, cytotoxicity of DOX in the treatment of GBM and epidermoid cancer spheroids and also demonstrate the significance of 3D cell culture models over 2D cells in drug delivery and discovery research.

Doxorubicin hydrochloride (DOX.H) is a naturally fluorescent anthracycline antibiotic that has been isolated from *Streptomyces peucetius var. caesius*. It’s a hydroxylated derivative of daunorubicin, a water-soluble anticancer agent. To verify the US-induced sonoporation in U-251 MG tumour spheres, the intracellular fluorescence intensity of DOX.H was evaluated by flow cytometry. DOX uptake was measured with and without US after 48 h of incubation by using U-251 MG (Figure 8A) and A431 (Figure 8B) tumour spheres. Sonoporation is the development of pores in cell membranes produced by US [9]. According to estimates, the pore size may range from 1 nm to several micrometres [9]. Pore resealing occurs rapidly, typically ranging from a few seconds to up to 180 seconds in duration [9]. During this period, penetration of DOX into tumour spheres was observed. As seen in Figure 8A, the DOX uptake in U-251 MG tumour spheres rose to almost 96.26 % and 97.88 % following 1 minute and 3 minute US treatments, respectively, compared to DOX without US (34.03 %). While in Figure 8B, it can be seen that, DOX uptake in A431 tumour spheres increased to almost 98.37 % and 99.69 % following 1 minute and 3 minute US treatments, respectively, compared to DOX without US (68.60 %). The percentage of tumour sphere cells, that encounter DOX is shown in Figure 8C and we observed DOX increase with US due to the drug’s migration through the tumour sphere. Drug diffusion in 3D model is enhanced by US (as indicated by % uptake). Figure 8D depicts the average quantity of DOX entering the cells in a tumour sphere. The higher mean fluorescence index values with US treatments indicate that sonoporation has occurred in the cells. Ultimately, it demonstrates that the US is increasing both the percentage of cells encountering DOX and the amount of DOX going into each cell. This leads to improved cytotoxicity in the 3D culture model with US exposure, as we proved in the previous section (Figure 7). According to our findings, transient DOX sonoporation augmented by ultrasonic therapy in 3D tumour spheres is a promising strategy for enhancing therapeutic efficacy and selective cytotoxicity in the future.

**Figure 8:**
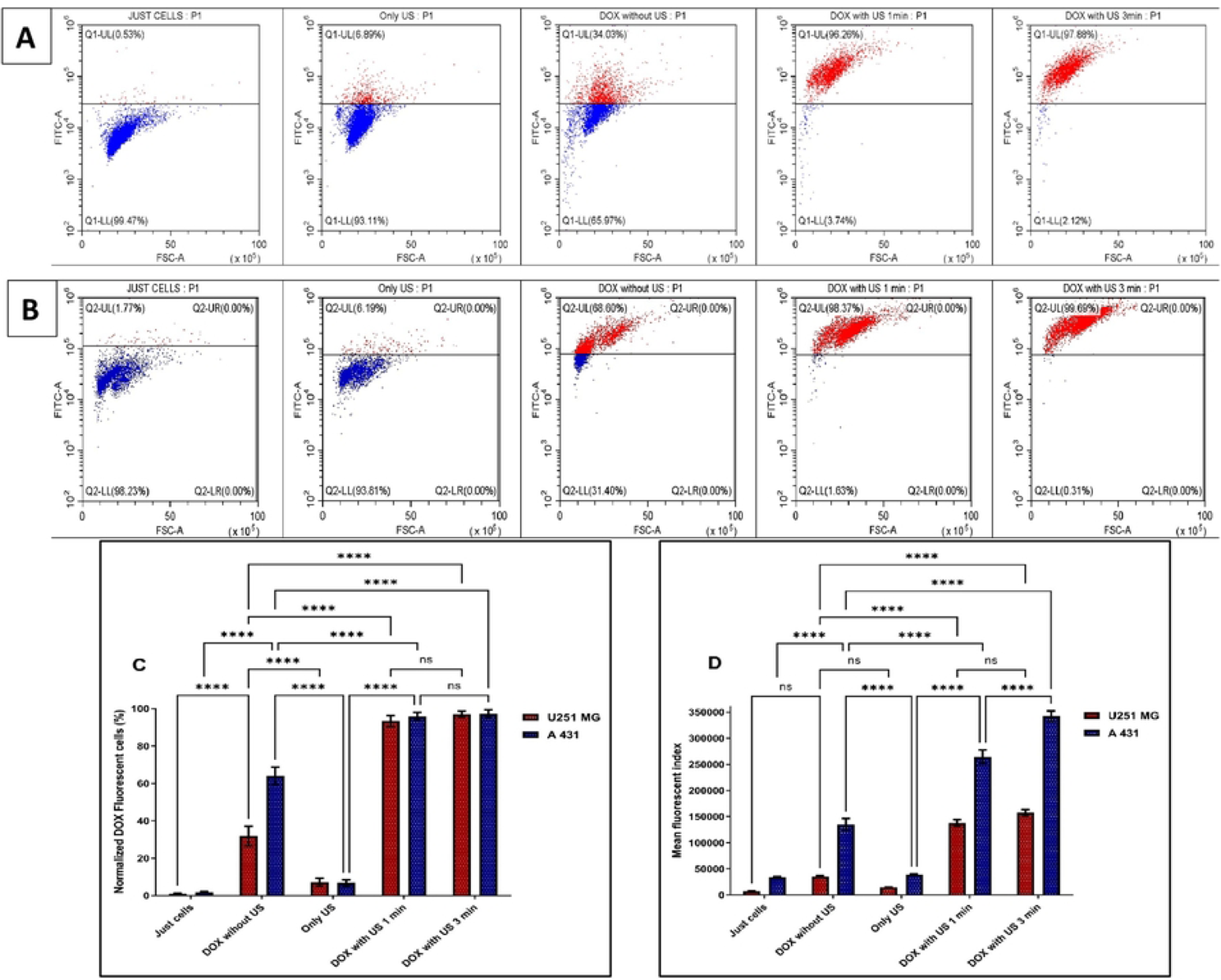
Ultrasound-enhanced penetration of DOX.H fluorescence analysis using flow cytometry of U-251 MG and A431 tumour spheres. A) U-251 MG B) A431 C) The normalized % of DOX positive cells was then measured at 48 h post treatment incubation and represented as a bar chart. All the data points were statistically significant except DOX with US 1 min and 3 min treatment times. ns, not significant, ****p ≤ 0.0001 D) The mean fluorescence index was measured at 48 h post treatment incubation, used to plot the values on columns to identify the amount of DOX getting into tumour sphere. All the data points were statistically significant except U-251MG DOX with US 1 min and 3 min treatment times. ns, not significant, ****p ≤ 0.0001

## Conclusion

US treatment can efficiently trigger cell death in 3D GBM and epidermoid carcinoma tumour spheres, a process contingent on time, dosage, and treatment frequency. Additionally, US has the capacity to diminish both 3D GBM spheroid growth and cellular proliferation while inducing damage to the tumour microenvironment. Our findings highlight crucial constraints in the prospective clinical application of US, indicating that an approach involving multiple treatments is more advantageous compared to a single treatment. Our findings illustrate the significance of US exposure time adjustment in order to achieve cytotoxicity and/or effective drug delivery. Less than three minutes of US exposure duration is optimal for transient sonoporation without damaging tumour spheres. When US exposure duration exceeds 3 minutes and/or treatment frequency is increased, enhanced cytotoxicity, persistent pore formation in membranes, lysis, subsequent leakage of cellular content, cell inactivation, or cell death occur in the tumour sphere.

The combination of US and TMZ enhanced the efficiency of GBM and epidermoid carcinoma treatment by enhancing TMZ induced cytotoxicity in 3D tumour spheres compared to 2D cells. DOX, as a reporter, revealed that US significantly enhances drug diffusion within 3D models and facilitates drug uptake into cells within tumor spheres. This leads to improved cytotoxicity in the 3D culture models with US, which is not evident in the 2D culture model, in which the cells are bathed in drug and the effects of sonoporation are muted. In conclusion, our study underscores the importance of utilizing 3D cell culture models in preclinical research. Traditional approaches involving 2D cell culture, followed by animal testing and clinical trials, have led to a high rate of clinical trial failures up to 95% due to their inability to accurately predict human efficacy and toxicity. Employing 3D cell culture models offers a more relevant and reliable platform for preclinical assessments, potentially improving the success rate of future clinical trials.

## Acknowledgments

This study was supported by Irish Research Council, Government of Ireland Postdoctoral Fellowship 2023, Project ID: GOIPD/2023/1288; Science Foundation Ireland (SFI) under Grant Number 17/CDA/4653 and 21/FFP-A/9189, and funded through Teagasc Walsh Fellowship. Authors also thank ESHI Research Institute and Focas Research Institute in TU Dublin for the use of facilities and support of technical staff.

## Data availability statement

All relevant datasets that support the findings of this study are uploaded into OSF and can be accessed using the following DOI link. (We have uploaded all datasets to OSF and we will provide relevant DOI link prior to publication)

## Ethical approval

The research project was approved by TU Dublin Research Ethics and Integrity Committee (approval granted 1^st^ March 2019) and involved the use of human samples. All experiments in the present study were conducted under ethical approval, between the dates of January 2021 and January 2023. Human cancer cell lines (U-251 MG and A431) were used in the study. These are cell lines obtained from reputable commercial cell banks, these are established, commercially available cell lines and consent was not obtained from the original donors. Animal tissue (fetal calf serum) was also used in the study. This was obtained from a reputable commercial company.

## Informed consent

For this type of study, formal consent is not required.

